# Sensorimotor adaptation to altered postural dynamics implemented via closed-loop neuromuscular electrical stimulation

**DOI:** 10.64898/2026.02.14.705802

**Authors:** Ryosuke Murai, Shota Hagio, Daichi Nozaki

**Author notes:** Corresponding author: Ryosuke Murai, Center for Information and Neural Networks (CiNet), National Institute of Information and Communications Technology, Osaka 565-0871, Japan, Daichi Nozaki, Graduate School of Education, The University of Tokyo, Tokyo 113-0033, Japan.

## Abstract

Studying sensorimotor adaptation in whole-body motor tasks such as locomotion and postural control remains challenging because well-controlled mechanical perturbations typically require large, specialized apparatus that constrains natural movement. Here, we introduced a novel perturbation system that alters musculoskeletal dynamics using closed-loop neuromuscular electrical stimulation (NMES) and examined how the human postural control system adapts to these altered dynamics during quiet standing. By applying NMES to the tibialis anterior as a function of anterior–posterior body sway, we imposed artificial postural dynamics. Analyses of postural sway revealed robust, systematic adaptation, with distinct patterns across perturbation types. These findings demonstrate that closed-loop NMES can impose controllable, movement-specific dynamics without mechanical constraints, while also revealing the adaptability of human postural control to externally imposed sway dynamics.

## Introduction

Human bipedal postural control serves as the foundation for a wide range of daily activities, from standing and walking to more complex motor tasks. To maintain the mechanically unstable upright stance, the central nervous system continuously integrates multimodal sensory inputs such as visual, vestibular, and somatosensory information [Fitzpatrick & McCloskey, 1994; Horak, 2006; Peterka, 2002; Oie et al., 2002, Maurer et al., 2006]. In daily life, the postural control system must also adapt to changes in both internal and external environments: standing on a compliant surface requires a different postural control strategy to standing on a rigid surface; changes in musculoskeletal properties due to growth, aging or muscle fatigue demand adjustments in motor commands to maintain standing stability.

Such a sensorimotor adaptation process of the postural control system has often been studied by investigating adaptations to transient or sustained external perturbations. These perturbations include muscle/tendon vibration [Fransson et al., 2000; Tjernström et al., 2002], changes in support surface properties [Patel et al., 2008; Patel et al., 2011], rotating or moving platforms [Dietz et al., 1993; Polastri et al., 2012; Fujiwara et al., 2012], and kinematic constraints by exoskeletons [Ringhof et al., 2019; Fasola et al., 2019], which were typically delivered in a constant or pseudo-random manner.

In the field of motor control research, ‘dynamics modulation’ approach has also been a well-established approach to investigate sensorimotor adaptation processes. In this approach, dynamics of the body during movement are systematically manipulated by an external perturbation, and the adaptation process to the novel dynamics generated by the perturbation is examined. This paradigm has made it possible to obtain detailed information about the adaptation process, compared to a simple open-loop perturbation.

One of the most representative studies using this approach is one by Shadmehr and Mussa-Ivaldi [Shadmehr & Mussa-Ivaldi, 1994], where participants performed repeated arm reaching movements while grasping the handle of a robot manipulandum. The manipulandum changed the limb’s dynamics by producing force depending on the hand velocity, and the adaptation to the novel dynamics was observed. While this paradigm has been used mainly for simple movements like arm reaching [Shadmehr & Mussa-Ivaldi, 1994; Lackner & Dizio, 1994; Sainburg et al., 1999; Thoroughman & Shadmehr, 2000], a similar strategy can also be applied to movements with a larger degree of freedom. For example, Cajigas and colleagues used a robotic system to study motor adaptation of gait patterns through motion-based mechanical perturbations during successive gait cycles [Cajigas et al., 2017]. While such a dynamics manipulation by mechanical perturbation is a powerful tool for studying sensorimotor adaptation, it generally requires large-scale equipment and its interference with bodily movement is inevitable, especially for whole-body tasks such as postural control.

To address this limitation, we propose an alternative approach using neuromuscular electrical stimulation (NMES), a technique that elicits involuntary muscle contractions through electrical pulses delivered via skin-surface electrodes. Traditionally, NMES has been used in rehabilitation, sports medicine, and physical therapy to improve muscle strength [Babault et al., 2007; Billot et al., 2010], reduce muscle atrophy [Baldi et al., 1998; Dirks et al., 2014], and restore motor function [Yan et al., 2005; Ajiboye et al., 2017]. In contrast to these conventional uses, this study aimed to alter body dynamics by modulating the NMES pattern (e.g., intensity and timing) based on the kinematics (i.e., closed-loop NMES). In other words, we sought to use muscles themselves as natural perturbing devices. Given that NMES requires no large-scale equipment and can induce contractions in various muscles simply by changing stimulation sites, NMES-based perturbation represents a promising approach to investigating sensorimotor adaptation across a wide range of movements without interfering with natural motion.

In the present study, we examined sensorimotor adaptation to altered postural dynamics during quiet standing, using a novel perturbation system based on a closed-loop NMES. Specifically, we applied amplitude-modulated NMES to the tibialis anterior (TA) muscle in response to body sway, thereby altering postural control dynamics in the sagittal plane. We selected the TA as the stimulation target because it typically exhibits relatively low tonic activity during quiet standing and is recruited mainly during transient corrective responses; thus, the stimulation could be treated primarily as an external perturbation rather than as a direct augmentation of the muscles that primarily contribute to tonic postural stabilization. Using this system, we investigated adaptation responses to two types of perturbations: position-dependent and velocity-dependent stimulation patterns.

## Results

### Alteration of postural dynamics by a closed-loop NMES system

We briefly introduce a closed-loop NMES perturbation system that can alter postural dynamics. Assuming the body during upright stance as a single inverted pendulum rotating within the sagittal plane (Fig. 1a), the equation of motion of the body in the anteroposterior (AP) direction can be written as:

**Fig. 1.**
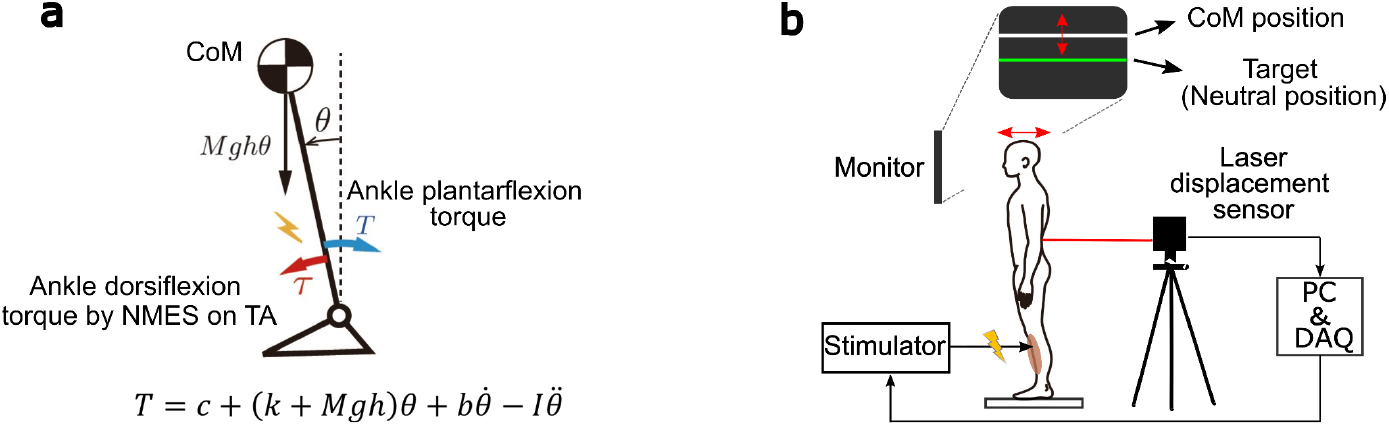
Manipulation of the postural control dynamics using closed-loop NMES. (a) Concepts of the postural control perturbation system with neuromuscular electrical stimulation (NMES) on tibialis anterior (TA). NMES intensity was modulated based on the CoM position and velocity in the AP direction. (b) Experimental setting of the quiet standing task. Participants kept queit standing on a 40 × 60 cm flat platform. NMES, either constant, step-like, or sway-dependent, were applied on TA based on the perturbation schedule (see f). A monitor in front of a participant’s head displayed the real-time CoM position (white line) and the neutral position (green line) on a predetermined schedule. Participants were required to keep their CoM close to the neutral position when the feedback was given.

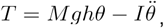

where *T* is the ankle plantarflexion torque, *θ* is the angular displacement from the vertical line, and *M, H, I*, and *g* are constants representing body mass, center of mass (CoM) height, moment of inertia around the ankle, and gravitational acceleration, respectively. The *θ* is assumed to be small enough for to sin*θ* ≈ *θ* hold.

Let *τ* be the externally applied perturbation torque to induce ankle dorsiflexion, and if *τ* can be modulated based on the position and velocity of the CoM in the AP direction (i.e., 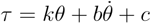, where *k, b*, and *c* are constants), then the resulting equation of motion is

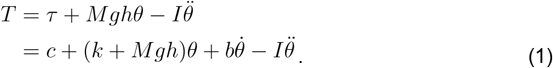

This equation indicates that novel postural control dynamics can be created by varying the value of the constants *k, b*, and *c*.

### Identification of NMES-torque relationship

Instead of using mechanical devices such as an exoskeleton or a wired device [Khan et al., 2019; Luna et al., 2020; Potocanac et al., 2017], we conceived the idea of employing NMES to implement closed-loop perturbation. Specifically, ankle joint torque perturbations [i.e., *τ* in Eq. (1)] were generated by modulating NMES intensity based on AP body sway, which was measured using a laser displacement sensor (Fig. 1b; see *Methods*). We assumed that the transfer function, which converts the NMES into the ankle joint torque, could be described as a first-order system [Rouhani et al., 2016]: *G*(*s*) = *K* / (1 + *as*), where *K* and *a* are constants.

We measured the ankle joint torque in response to NMES applied to the TA using biphasic rectangular pulses at 20 Hz, with the stimulation amplitude sinusoidally modulated at frequencies of 0.07, 0.15, 0.3, 0.75, 1.2, 1.5, and 2 Hz (Fig. 2a, b). Figure 2c shows the Bode diagram illustrating the gain and phase of the ankle joint torque relative to the frequency of NMES amplitude modulation. Based on this result, we obtained *K* and *a* of the transfer function for each participant. Notably, while the identified system describes how NMES is converted into ankle joint torque, what is required here is to determine the appropriate NMES input required to realize a desired ankle joint torque pattern. We achieved this using the inverse transfer function as *G*^−1^(*s*) = (1 + *as*)/{( 1 + *as*)/*d* + *K*}, where we set *d* = 1000, instead of using the simple inverse transfer function (i.e., *G*^−1^(*s*) = (1 + *as*)/*K*). We introduced this correction to reduce the simple inverse transfer function’s susceptibility to high-frequency noise, an approach analogous to replacing an ideal derivative with a dirty (filtered) derivative in PID control [Li & Chong, 2006].

**Fig. 2.**
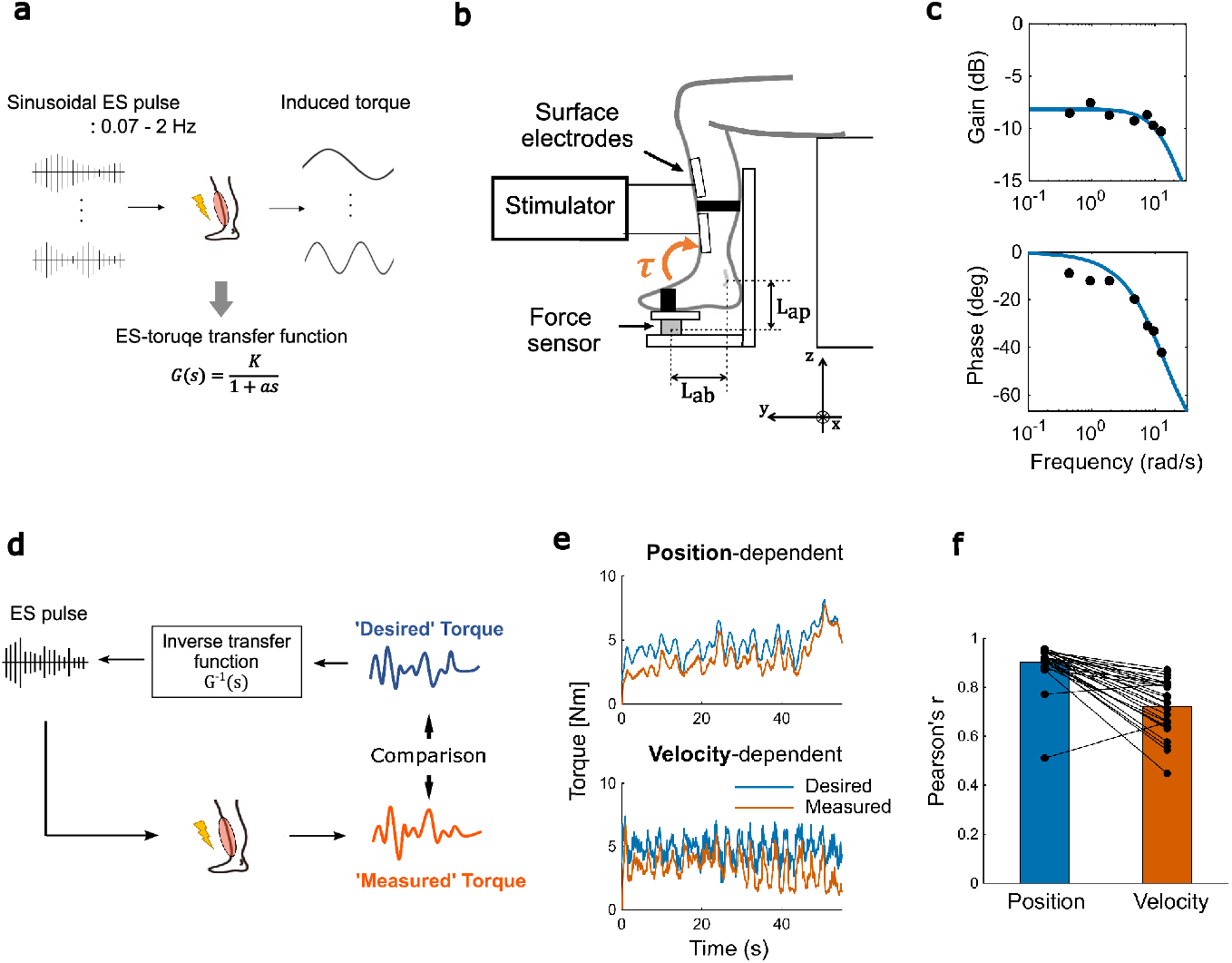
Experiment 1: Estimation of ES-torque relationship. (a) Schematic illustration of the estimation of the ES-torque transfer function. (b) Torque measurement system. Ankle dorsiflexion torque during NMES application on TA was bilaterally measured by the 6-axis force sensor mounted under the feet. The feet and shank were secured to the device using hook-and-loop fasteners. (c) Bode diagrams of the ES-torque transfer function estimated from sinusoidal NMES input with seven different frequencies for the right leg of a representative participant. Dots represent data from each frequency. (d) Schematic illustration of the evaluation of the accuracy of the ES-torque transfer function. (e) Time series of ‘desired’ (blue) and ‘measured’ (red) torque for the right leg of the same participant. (f) Mean (n = 28) Pearson’s r between ‘desired’ and ‘measured’ torque for the position- and velocity-dependent condition. Dots represent individual data. The mean values were computed by first applying Fisher’s z-transformation to each r, averaging the z-values, and then applying the inverse transformation.

### Validation of the NMES-torque relationship

To evaluate whether the proposed approach could generate a desired temporal pattern of ankle joint torque, we conducted a validation experiment using two types of ‘desired’ torque profiles: one position-dependent and the other velocity-dependent. The desired torque profiles were generated by multiplying a constant gain with either the position or velocity of the AP sway of CoM, obtained from data recorded beforehand during 60 seconds of quiet standing. The NMES patterns required to produce each desired torque profile were then estimated using the inverse ES–torque transfer function (Fig. 2d). These NMES patterns were then applied to the participants, and the resulting ankle joint torque was directly measured to assess how accurately it reproduced the corresponding desired torque profile.

Figure 2e shows a representative example comparing the temporal profiles of the desired and measured ankle joint torque. Although minor deviations were observed, the measured torque closely followed the desired torque. Figure 2f summarizes the strength of this relationship, showing significant correlations between the desired and measured torque in both torque profiles. Specifically, Fisher’s *z*-transformed Pearson’s correlations were *r* = 1.57 ± 0.05 for the position-dependent torque profile, and 0.94 ± 0.04 for the velocity-dependent torque profile (corresponding to *r* = 0.92 [95% CI: 0.90–0.93] and *r* = 0.74 [95% CI: 0.70–0.77], respectively). These results suggest that the proposed approach can reasonably generate perturbation torque at the ankle joint [i.e., *τ* in Eq. (1)].

### Adaptation to closed-loop perturbations

We then applied ankle joint torque perturbations based on either the CoM position (position-dependent) or CoM velocity (velocity-dependent), using the closed-loop NMES system while participants maintained quiet standing for 30 seconds. Following this period, a step-like NMES perturbation was introduced to evaluate the postural control system’s response (i.e., step response). This sequence (sway-dependent perturbation followed by a step-like perturbation) was repeated 12 times, and followed by a 60 sec washout phase during which NMES was completely turned off (Fig. 3a). Figure 3b shows the time course of AP sway during quiet standing under the velocity-dependent condition for a representative participant.

**Fig. 3.**
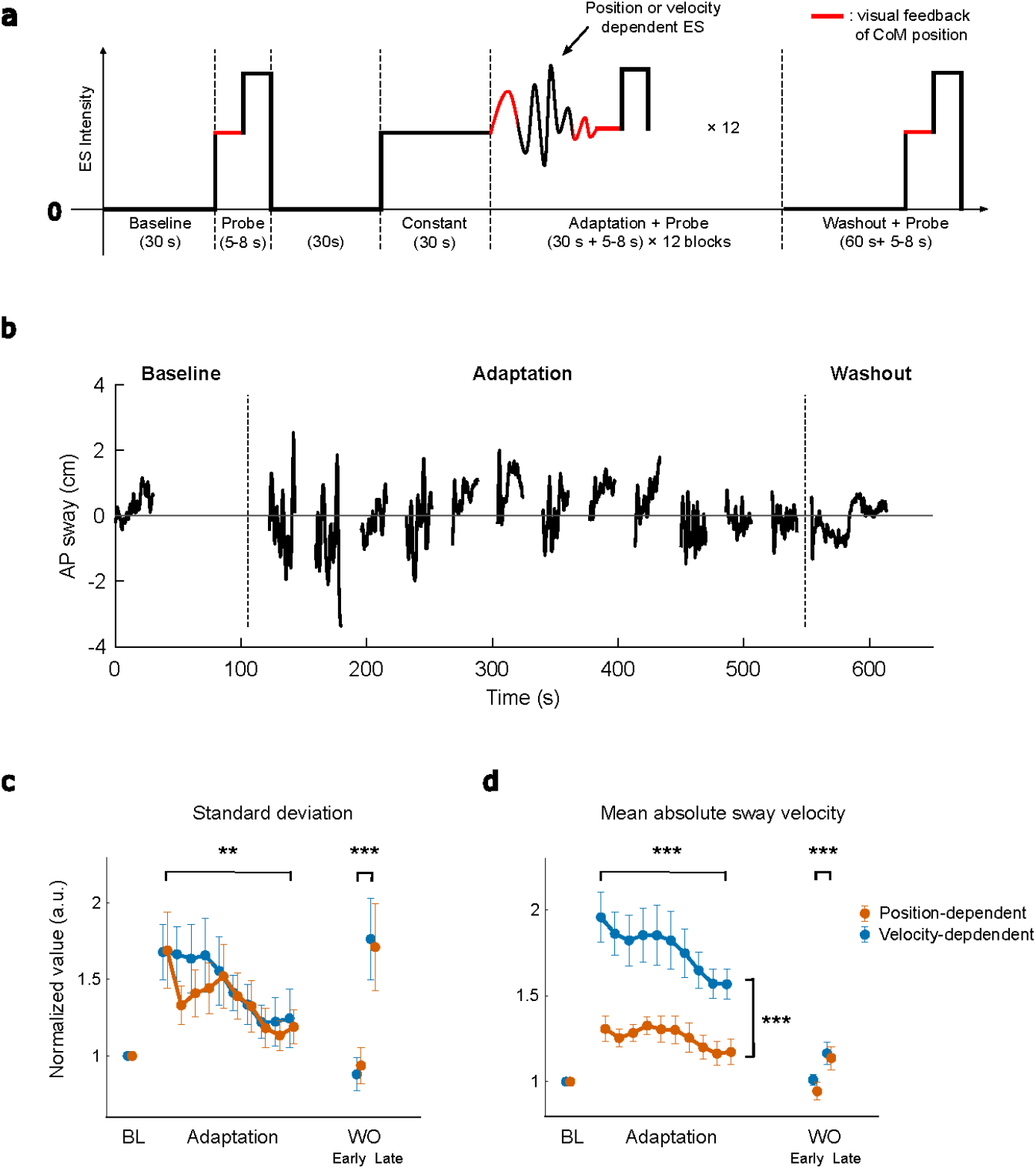
Experiment 1: Changes in AP sway in the quiet standing task. (a) Perturbation schedule of the quiet standing task. The red lines correspond to the time when the real-time feedback of the CoM position was provided on the monitor (see Fig. 1b). (b) Time course of AP sway for a representative participant during the quiet standing task (velocity-dependent condition). Only the no-visual-feedback segments of the sway-dependent ES parts during the adaptation phase, as well as the baseline and washout phases, are shown; probe parts are excluded. Time is aligned to task onset; an AP sway of 0 cm corresponds to the predetermined neutral position. (c, d) The change in the mean values of two behavioral measures for the position-dependent (n = 14) and velocity-dependent (n = 11) conditions in Experiment 1: (c) standard deviation of CoM sway and (d) mean absolute CoM sway velocity. Each value was individually normalized by the baseline value before being averaged across participants, and then moving-averaged with a window width of three blocks. Error bars mean standard error of the mean (SEM) across participants. The asterisk indicates significant main effects of time point (difference between the first and last time point in the adaptation phase or between the early and late washout)) or condition (difference between the position- and velocity-dependent conditions) (**: p < 0.01; ***: p < 0.001). BL, baseline; WO, washout.

Figure 3c, d illustrate changes in two behavioral measures during the quiet standing task. As shown in Figure 3c, the standard deviation of the AP sway increased during the early adaptation phase due to the perturbation but gradually decreased in both perturbation conditions. A linear mixed-effects model revealed a significant fixed effect of time point (F(1, 23.001) = 12.182, p = 0.002), but not of condition (F(1, 8.560) = 0.108, p = 0.750) or the time point * condition interaction (F(1, 23.001) = 0.062, p = 0.806). For the mean absolute velocity, however, significant fixed effects were found not only for time point (F(1, 23.000) = 20.105, p < .001), but also for condition (F(1, 10.460) = 24.609, p < .001) and their interaction (F(1, 23.000) = 4.751, p = 0.040) (Fig. 3d). A simple main effects test revealed that the decrease across time points was significant in the velocity-dependent condition (t(10) = 4.989, p < .001), but not in the position-dependent condition (t(13) = 1.615, p = 0.130). This difference may reflect the distinct nature of the two perturbation types. In the velocity-dependent condition, a reduction in sway velocity directly reduced NMES intensity thereby contributing to the stabilization of upright posture. In contrast, in the position-dependent condition, reducing sway velocity did not necessarily decrease the sway magnitude, limiting its stabilizing effect.

In the washout phase, these parameters initially returned to baseline levels (Fig. 3c, d WO early), but subsequently increased above baseline in the late washout phase (Fig. 3c, d WO late), with a significant fixed effect of time point for the standard deviation (F(1, 10.340) = 17.711, p = 0.002) and mean absolute velocity (F(1, 13.023) = 16.768, p = 0.001).

### Stabilogram diffusion analysis

While the behavioral parameters examined above are widely used metrics that quantify the spatiotemporal characteristics of postural sway, they can be influenced by non-oscillatory components such as drift of the mean CoM position. Thus, in addition to the analysis based on the behavioral measures, we conducted stabilogram diffusion analysis (SDA; [Collins & De Luca, 1993]). SDA has been widely used to evaluate the postural dynamics governed by short-term (open-loop) and long-term (closed-loop) control mechanisms (Fig. 4a). We speculated that SDA could more robustly capture the characteristic adaptations to both types of sway-dependent perturbations.

**Fig. 4.**
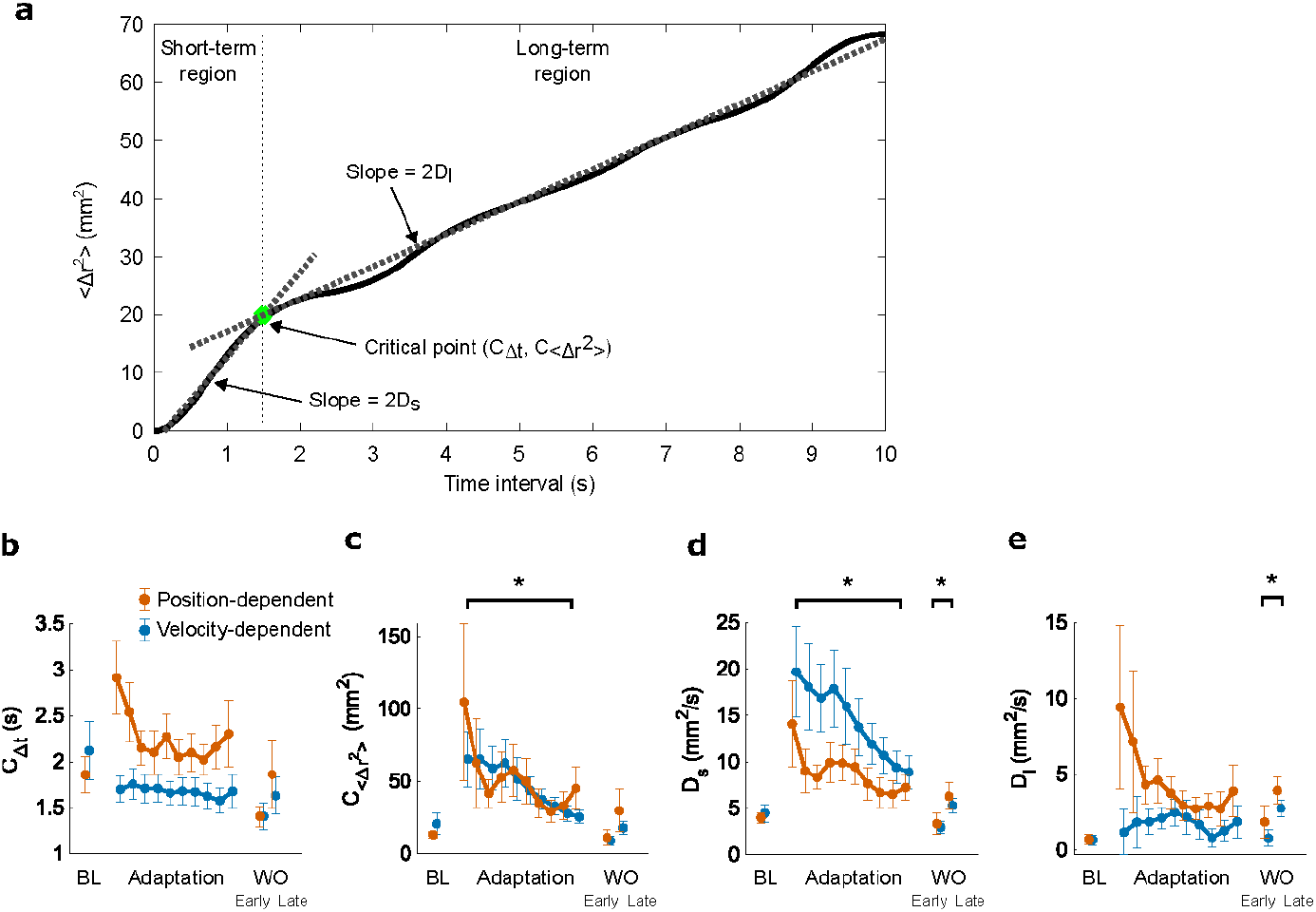
Experiment 1: SDA results of the quiet standing task. (a) Example of SDA plot and calculation of SDA parameters. (b–e) Change in the mean values of four SDA parameters for the position- and velocity-dependent conditions in Experiment 1: critical point coordinates (b) *C*_Δ*t*_ and (c) 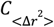 and short- and long-term diffusion coefficients (d) *D*_*s*_ and (e) *D*_*l*_. Error bars means standard error of the mean (SEM) across participants. The asterisk indicates significant main effects of time point (difference between the first and last time point in the adaptation phase or between the early and late washout) (*: p < .05).

Figure 4b–e illustrate the changes in the four SDA parameters in the baseline, adaptation and washout phase for both conditions. The value 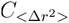 of and *D*_*s*_ showed an initial increase immediately after the perturbation onset, followed by a gradual decrease along adaptation for both types of perturbations (Fig. 4c, d). The significant fixed effects were found for time point (F(1, 30.865) = 5.226, p = 0.029 and F(1, 22.999) = 7.873, p = 0.010 for 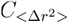 and *D*_*s*_, respectively), but not for condition (F(1, 32.210) = 0.169, p = 0.684 and F(1, 4.950) = 0.022, p = 0.888) or time point * condition interaction (F(1, 30.865) = 0.640, p = 0.430 and F(1, 22.999) = 0.003, p = 0.957). For the *C*_Δ*t*_ and *D*_*l*_, however, the decrease through the adaptation was found only for the position-dependent condition (Fig. 4b, e), although no significant fixed effects were found for time point * condition interaction (F(1, 31.125) = 1.124, p = 0.297 and F(1, 19.615) = 0.029, p = 0.866) for *C*_Δ*t*_ and *D*_*l*_, respectively.

In the washout phase, we found a significant fixed effect of time point for *D*_*s*_ (F(1, 11.536) = 8.705, p = 0.013) and *D*_*l*_ (F(1, 33.937) = 7.416, p = 0.010), but not for *C*_Δ*t*_ (F(1, 10.844) = 2.109, p = 0.175) or 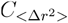 (F(1, 12.334) = 2.116, p = 0.171). These results indicate that the increase in sway during the washout phase was also captured in the SDA parameters (*D*_*s*_ and *D*_*l*_), in addition to the behavioral measures.

### Change in step response by the adaptation

We then examined how adaptation to altered postural dynamics affected compensatory responses to transient step-like perturbations. Qualitatively, we observed two key features in the step response during the adaptation phase (Fig. 5a, b). First, the CoM sway following the step-like perturbation exhibited a distinct pattern across the two perturbation conditions. Compared to the velocity-dependent condition, the CoM in the position-dependent condition tended to return to its neutral position more rapidly after the peak of the step response. This was supported by a smaller area under the curve (AUC) of the CoM sway from the response peak to 1 second afterward—reflecting the magnitude of backward CoM displacement—and by a greater negative sway velocity from 1 second onward (Fig. 5a–c). In a linear mixed-effects model, change in AUC showed a significant fixed effect of perturbation condition (F(1, 11.091) = 6.147, p = 0.030), but not of time point (F(9, 207.000) = 0.509, p = 0.867) or the time point * condition interaction (F(9, 207.000) = 0.980, p = 0.458) (Fig. 5c), suggesting that this difference arose from the very early part of the adaptation phase. Second, the peak magnitude of the subject-averaged step response did not monotonically decrease with adaptation but rather fluctuated, particularly exhibiting an inverted U-shape change in the velocity-dependent condition (Fig. 5a, d). This pattern suggests that the overall adaptive strategy involved not only increasing body stiffness via muscle co-contraction but also changes in the control policy itself because an increase in stiffness alone was unlikely to raise the peak magnitude.

**Fig. 5.**
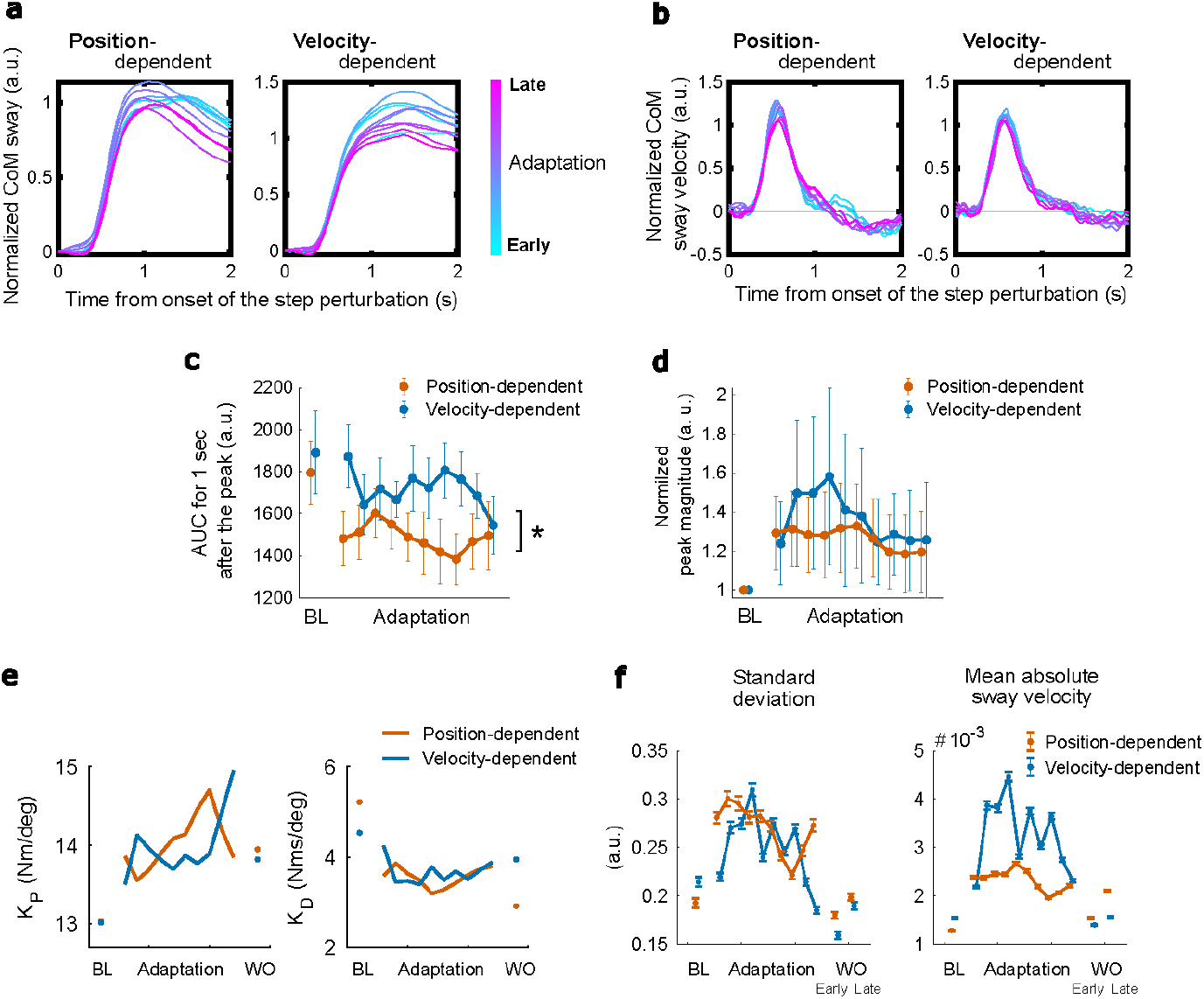
Experiment 1: Step response in the quiet standing task. Change in (a) position and (b) velocity profiles of the step responses in the adaptation phase. Step responses were individually normalized by the peak magnitude of the baseline phase, moving-averaged with a window width of three blocks, and then averaged across participants. The color of the lines changes from cyan to magenta as the task proceeds. (c) Area under the curve (AUC) and (d) peak magnitude of the step response. The peak magnitude was individually normalized by the baseline value. Error bars mean SEM across participants. The asterisk indicates significant main effects of condition (*: p < .05). (e) Estimation of proportional (left) and derivative (right) gains during the quiet standing task using a one-dimensional inverted-pendulum with a PID controller. Gains were estimated to best reproduce the step responses in the quiet standing task. The step responses were subject-averaged for all phases, and additionally moving-averaged only in the adaptation phase before the estimation. (f) Simulation of the 40-seconds quiet standing task using the PID controller with the estimated gains in (e) could reproduce the changes in the behavioral measures in Experiment 1. The simulation was iterated 100 times for each set of gains. Error bars mean SEM across iterations.

By comparing the subject-averaged step responses with simulations from a postural control model with a PID controller, we estimated changes in postural control (proportional and derivative) gains during the quiet standing task. The results of the estimation suggested that these gains were flexibly modulated during the adaptation phase and did not return to their baseline levels within the 1-min washout period (Fig. 5e). In addition, using the same PID-based postural control model with the estimated gains, we simulated postural sway during the baseline, adaptation, and washout phases and computed the behavioral measures. These gain estimates, which rely on step responses measured only once at each time point and should therefore be regarded as approximate, yielded simulations that qualitatively reproduced the characteristic adaptation pattern during the adaptation phase, including the distinct patterns of change in mean absolute velocity between perturbation types, although it produced unexpectedly low values in the first block of the velocity-dependent condition due to the high derivative gain (Fig. 5f). In contrast, during the washout phase, the changes did not show an increase beyond the baseline level as observed in the experimental data. Instead, they tended to decrease in the early washout and return to baseline in the late washout. Altogether, these results suggest that the postural control system adapted to the perturbation by modulating postural control gains.

### Effects of muscle fatigue

Although the frequency of NMES used in this study (20 Hz) is generally associated with lower levels of muscle fatigue—that is, less reduction in force production—compared to higher frequencies [Moritani et al., 1985], the prolonged application of NMES (∼10 minutes per set) may still have induced some degree of muscle fatigue. To assess this possibility, we examined the potential effects of muscle fatigue on our results. Specifically, we compared ankle joint torque produced by NMES before (pre) and after (post) the quiet standing task. Figure 6a shows a representative participant’s ‘measured’ torque before and after the task. While the correlation between ‘desired’ and ‘measured’ torque remained high even after the completion of the task (Fig. 6b, Fisher’s *z*-transformed Pearson’s *r* was 1.01 ± 0.05, corresponding to *r* = 0.77 [95% CI: 0.72–0.80]; the difference between pre and post was not significant: t(27) = -2.0311, p = 0.0522), mean torque production significantly declined by 44% (Fig. 6c, t(27) = 11.7312, p < .001). Thus, the muscle fatigue did occur due to the prolonged NMES application.

**Fig. 6.**
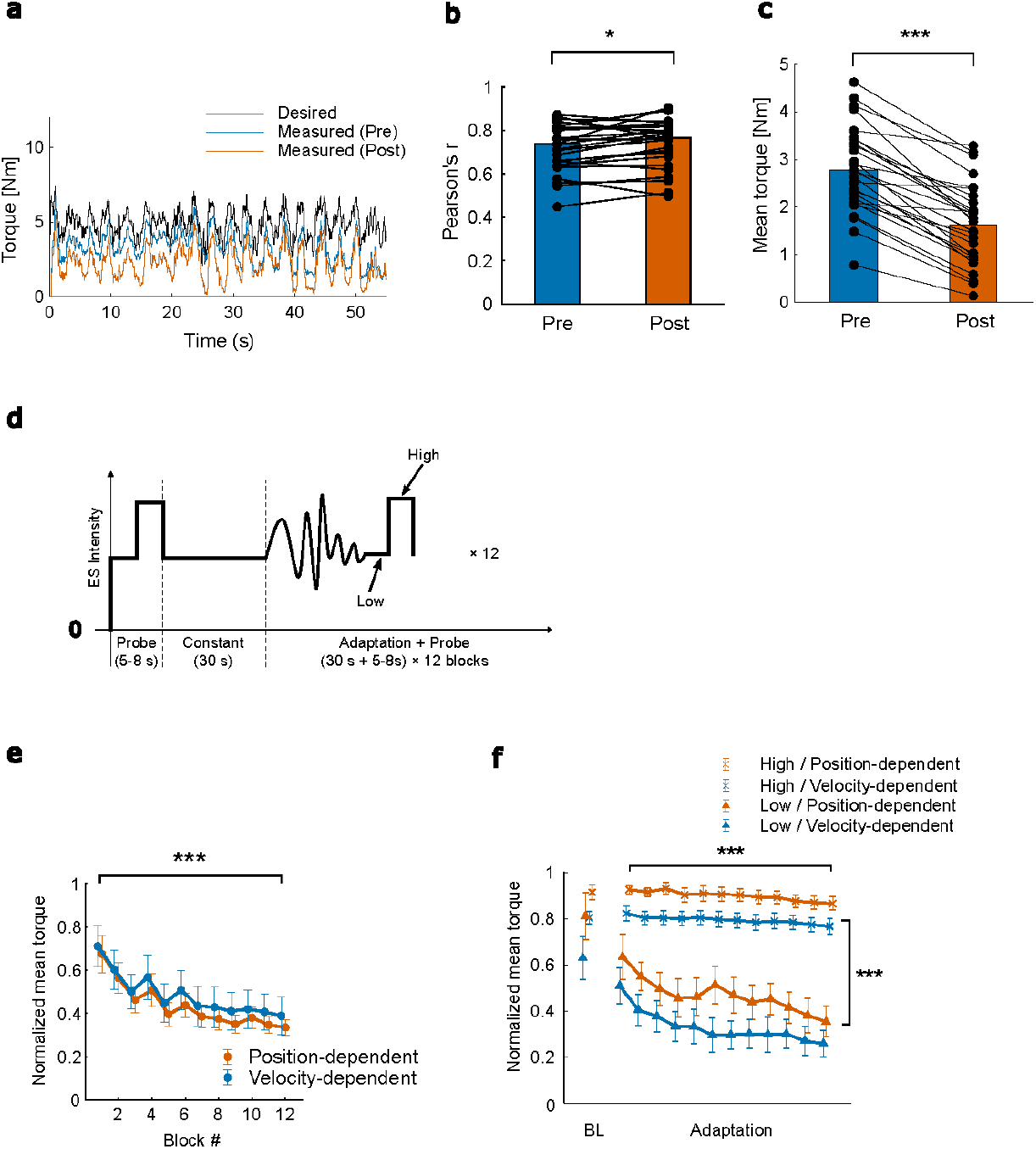
Effects of muscle fatigue on perturbation torque production. (a–c) Comparison of the torque induced by the pre-designed velocity-dependent NMES before (pre) and after (post) the quiet standing task in Experiment 1. (a) Time series of ‘desired’ (black) and ‘measured’ (before [blue] and after [red] the quiet standing task) torque for the right leg of a representative participant. Although the correlation between ‘desired’ and ‘measured’ torque remained high (b; n= 28), torque production significantly decreased after the prolonged NMES application in the quiet standing task (c). Dots represent individual data. The asterisks indicate statistical significance (*: p < .05; ***: p < .001). (d–f) Experiment 2: Real-time measurement of torque induced by NMES similar to those used in the quiet standing task. (d) Perturbation schedule in the real-time torque measurement. ES intensity was predetermined to replicate the one in the queit standing task in Experiment 1. (e) Change in the mean torque induced by pseudo sway-dependent NMES in the adaptation phase. Mean torque was normalized by ‘reference’ torque, and then averaged across individual legs (n = 8 from 4 participants). (f) The same procedure was performed to calculate the change in the mean torque induced by the low and high intensities of step-like NMES. Mean torque was normalized by ‘reference’ torque, and then averaged across individual legs (n = 8 from 4 participants). Mean and standard deviation were calculated for each block of the adaptation phase. Error bars correspond to SEM across individual legs. The asterisks indicate significant main effects of the time point or intensity (**: p < .01; ***: p < .001).

We further examined directly the influence of prolonged NMES on the torque production. In a subsequent experiment (Experiment 2; see *Methods*), ankle dorsiflexion torque was measured while participants were seated during 10 minutes of continuous NMES application that imitated the perturbation modulating with the postural sway in the quiet standing task (i.e., position-dependent or velocity dependent) (Fig. 6d). In Experiment 2, we found a significant decline in the torque production for the pseudo sway-dependent part over the 10-minute NMES application (Fig. 6e; F(1, 6.999) = 35.081, p < .001 for time point). Notably, the majority of this decline occurred in the early phase, with little further reduction observed after the fifth block. This time course differed from that of postural sway magnitude, which decreased continuously throughout the adaptation phase (Fig. 3c, d).

We also examined torque induced by step-like NMES, implemented by switching stimulation intensity from the low to high level. Torque induced by both high and low intensity levels gradually declined, with the decline being faster and more pronounced in the low intensity (Fig. 6f). We found significant fixed effects for time point (F(1, 43.000) = 32.058, p < .001), intensity (F(1, 43.000) = 201.568, p < .001), and time point * intensity interaction (F(1, 43.000) = 13.107, p < .001). While significant simple main effects were found for both levels of the intensity (t(23) = -2.089, p = 0.048 and t(23) = -4.412, p < .001 for low and high intensities, respectively), the decline in torque from the first to the last block was relatively small for the high-intensity level, with a mean decrease of approximately 7%. This greater torque decline in the low-intensity region was likely to have increased the difference in torque induced by low- and high-intensity NMES, thereby potentially amplifying the effective perturbation magnitude.

As a supplementary analysis, we performed a subgroup analysis for the quiet standing task in Experiment 1 by classifying participants as fatigued or less fatigued based on the decline in torque production (the reduction in mean torque after the quiet standing task ≥40% vs. <40%). Overall, the behavioral measures and step-response profiles appeared broadly similar between the two subgroups in both conditions, with no obvious quantitative differences (Supplementary Fig. 1), suggesting that fatigue severity did not substantially influence the observed behavioral patterns.

## Discussion

In this study, we investigated how the postural control system adapts to novel postural dynamics induced by a newly developed NMES-based perturbation system. Our perturbation system created novel postural dynamics by applying NMES to TA in a closed-loop manner, based on the CoM AP sway. The present results indicate that the postural control system adapted to the novel dynamics to stabilize upright stance, with the adaptation depending on the characteristics of the perturbations. Importantly, the proposed system served as a practical platform for systematically perturbing whole-body movements. Because the same NMES setup can readily target different muscles without large-scale equipment, it provides a versatile and accessible approach for probing sensorimotor adaptation across a wide range of motor tasks.

### Adaptation to the novel postural control dynamics

Avoiding falling is one of the most important objectives of postural control. The decrease in behavioral measures observed during the quiet standing task, which all contributed to maintaining an upright stance, suggests that the postural control system suppressed the excessive body AP sway caused by the perturbation to lower the risk of falling (Fig. 3c, d, Fig. 4c, d). Notably, despite these reductions in behavioral measures, the peak magnitude of the step response did not exhibit a monotonic decline during the adaptation phase (Fig. 5a, d). Together with the results of postural gain estimation (Fig. 5e), this suggests that the postural control system stabilized the upright stance by flexibly modulating postural control gains, rather than monotonically increasing them. At the same time, the elevated derivative gain in the early phase of the velocity-dependent condition (Fig. 5e) suggests that the postural control system may have initially increased the derivative gain to buffer the abrupt perturbation, after which a more adaptive response emerged, a strategy similar to those reported in arm-reaching tasks [Franklin et al., 2003; Osu et al., 2003]. It should be noted, however, that because the step response was measured only once at each time point and thus highly variable, the step-response data, as well as gain estimates and simulations based on it, and the corresponding interpretation remain somewhat speculative.

We observed distinct adaptation patterns between position- and velocity-dependent perturbations. For the behavioral results, we found a significant adaptive decrease in the absolute mean sway velocity, only for the velocity-dependent perturbation (Fig. 3e). Since decreasing the AP sway velocity in the position-dependent condition does not necessarily result in the reduced perturbation torque, i.e., stabilization of upright posture, this difference suggests that the postural control system adapted adeptly to perturbations based on their specific properties. Additionally, we found a difference in the step response between the two perturbations, where the backward CoM movement after the peak was greater for the position-dependent condition compared to the velocity-dependent condition (Fig. 5c). This rapid return to the neutral position contributes to lowering the NMES intensity in the position-dependent condition, further supporting the occurrence of the efficient adaptation of the postural control system; however, the step response data, as noted above, should be interpreted with caution given the limited sampling and substantial variability.

The results of SDA may also reflect the different characteristics of the two perturbations. Both the short- and long-term diffusion coefficients (*D*_*s*_ and *D*_*l*_) in the position-dependent condition exhibited a temporary increase after the perturbation onset, followed by a gradual decrease along adaptation, whereas such a trend was observed only for *D*_*s*_ in the velocity-dependent condition (Fig. 4d, e). Based on the assumption that the open-loop control schemes are utilized over the short-term region [Collins & De Luca, 1993], this common gradual decrease of *D*_*s*_ along adaptation is associated with the reduction of CoM sway through open-loop control without feedback information (e.g., muscle co-contraction) in both conditions. On the other hand, a different scenario can be considered for the long-term region, which is thought to reflect closed-loop control [Collins & De Luca, 1993]. For instance, consider the situation where the body starts to lean forward. If the velocity-dependent perturbation is applied, the forward sway initially increases perturbation torque. Still, as the forward sway velocity begins to decrease, perturbation torque also decreases. In such a situation, the CoM spontaneously moves backward if the ankle plantar flexion torque produced by triceps surae remains constant. This means that the velocity-dependent perturbation itself acted as a negative feedback control system, and consequently the degree of increase in *D*_*l*_ was relatively small. In contrast, the position-dependent perturbation does not have such a function. Accordingly, *D*_*l*_ increased after the perturbation onset, and the postural control system adapted to it by strengthening the closed-loop control. Overall, these findings suggest that the postural control system could change its adaptation strategies according to the nature of the perturbation.

### Increased postural sway during the washout phase

Our results demonstrated an increase in the behavioral measures above baseline levels during the washout phase (Fig. 3c, d WO late). Typically, however, such a change in behavior after the removal of a perturbation, referred to as aftereffects, subside monotonically over time and behavior returns to the baseline level [Bastian, 2008]. One possible explanation for this discrepancy is task-related fatigue. Given the task demands, adaptation likely increased postural control gains; under non-fatigued conditions, this would be expected to reduce sway below baseline in the early washout phase, followed by a gradual return toward baseline in the late washout phase along de-adaptation. In the current experiment, however, participants were required to stand for approximately 10 minutes, and such sustained standing may have induced fatigue, increasing postural sway relative to a non-fatigued state and thereby elevating the behavioral measures and SDA parameters above baseline during washout. Consistent with this interpretation, the behavioral measures derived from simulations in which the effect of fatigue was not incorporated exhibited a more typical washout pattern, although the estimated gains themselves did not fully revert to their baseline levels.

### Potential effects of NMES-induced muscle fatigue

Since continuous electrical stimulation on a specific muscle induces muscle fatigue [Moritani et al., 1985; Tepavac & Schwirtlich,1997], one may argue that the stabilization of upright stance observed in the quiet standing task was merely due to the decline in perturbation torque caused by the prolonged application of NMES. Indeed, we found that torque production by NMES of the same intensities became smaller after the quiet standing task (Fig. 6c). However, we consider it unlikely that the NMES-induced muscle fatigue alone can fully account for our results.

Real-time torque measurement during the application of NMES (Experiment 2), similar to that used in the quiet standing task, showed a rapid torque decline shortly after the NMES onset, especially in the low torque region (Fig. 6e, f). The greater torque reduction in the low torque region would have expanded the difference between the torque induced by low and high intensities of the step-like NMES, resulting in an increased peak of the step response in the late adaptation phase if no adaptation had occurred. Indeed, during the quiet standing task, the peak of the step response showed a temporary increase in the early-mid part of the adaptation phase for the velocity-dependent condition (Fig. 5d). Nonetheless, the peak of the step response decreased afterward, suggesting the occurrence of adaptation. In addition, while the large part of the torque decline occurred in the early adaptation phase in the real-time torque measurement (Fig. 6e), the change in the behavioral measures during the quiet standing task occurred also in the late adaptation phase (Fig. 3c, d), indicating the presence of adaptation alongside the influence of muscle fatigue. Finally, a subgroup analysis in Experiment 1, in which participants were divided into two groups based on the degree of the torque decline, revealed no qualitative differences between the two groups regarding the change in the behavioral measures (Supplementary Fig. 1).

In summary, our results suggest that the stabilization of upright stance observed in the quiet standing task can be attributed, at least partly, to sensorimotor adaptation of the postural control system, even though NMES induced substantial muscle fatigue.

### Novel perturbation system using closed-loop NMES

The present perturbation system with closed-loop NMES allows for postural dynamics modulation in quiet standing. More generally, task-level input–output relationships can also be manipulated via visual perturbations that alter the mapping between motor commands and their sensory consequences. For example, visuomotor transformations are commonly used to study sensorimotor adaptation of arm reaching, where the displayed cursor movement is distorted from the actual hand trajectory according to a specific law [Krakauer et al., 2000; Pine et al., 1996]. However, visual perturbation is difficult to apply for systematic, task-relevant manipulations in postural control or other whole-body movements such as gait, despite its convenience to implement, because these movements do not have an obvious task goal related to vision (e.g., moving a cursor toward the displayed target in the arm-reaching movement). In this sense, the present system, which realizes dynamics modulation by means of mechanical perturbation without interference to motion, has a significant importance. Furthermore, our system can be easily combined with perturbations to other modalities, such as visual [Lishman & Lee, 1973; Bugnariu & Fung, 2007] and vestibular systems [Cenciarini & Peterka, 2006, Héroux et al., 2015]. This approach will help to address important issues in postural control including multisensory integration.

The perturbation system developed in this study is also applicable to movements other than bipedal standing. This system has some advantages over classic mechanical perturbation devices such as robot manipulanda [Shadmehr & Mussa-Ivaldi, 1994] and exoskeleton robots [Cajigas et al., 2017], including minimal motion interference and no need for large equipment. In addition, it can in principle be utilized to perturb various types of movement by changing stimulation sites and patterns. Hence, it has potential to study motor adaptation of practical movements with multiple degrees of freedom and complexity, which has been difficult to address with conventional mechanical perturbations.

Although this perturbation system is a useful method to investigate motor adaptation for various movements, there are some points to consider. First, time lags between NMES onset and torque production are inevitable due to electromechanical delay [Cavanagh & Comi, 1979]. Such a time lag has a significant impact, especially on quick movements (e.g., pitching). For these movements, ES intensity needs to be determined predictively based on movement trajectories. Another point is that NMES that induces muscle contraction is above the sensory threshold, and participants will be aware of it. This point is important because the awareness of stimulus will, to a certain extent, affect the explicit adaptation strategy [Taylor & Ivry, 2012; Mazzoni & Krakauer, 2006; Taylor et al., 2014].

In conclusion, we demonstrated that the postural control system could adapt to the novel dynamics created by closed-loop NMES, depending on the nature of modulated dynamics. Furthermore, the NMES-based perturbation system proposed in this study proved to be a promising way to investigate the motor adaptation mechanisms, potentially for a variety of movements, including those that cannot be addressed by traditional mechanical perturbation.

## Methods

### Participants

A total of 21 males participated in this study (23.2 ± 2.6 years, 171.9 ± 4.6 cm, 60.7 ± 6.4 kg). The participants had no musculoskeletal or neurological disorders that impair the maintenance of upright standing. Each participant gave written informed consent prior to the experiments. The Ethics Committee of the University of Tokyo reviewed and approved the experimental protocol that was in accordance with the Declaration of Helsinki.

### Apparatus

The experiments in this study consisted of two components: torque measurement and a quiet standing task. During torque measurement, participants sat on a chair with their knee and ankle joint angles at 90° (Fig. 2b). The participants’ feet and shank were secured to a custom-made torque measurement system that incorporated a 6-axis force sensor (LFM-A-3KN, Kyowa Electronic Instruments Co., Ltd., Japan) to measure the ankle dorsiflexion torque exerted by NMES on TA. For the quiet standing task, participants stood on a 40 × 60 cm flat platform, and their body sway in the AP direction was measured by a laser displacement sensor (IL-1000, KEYENCE CORPORATION., Japan) aimed at the center of their waist (Fig. 1b). This body sway was considered as CoM sway in this study. A LCD monitor placed at a distance of about 80 cm in front of the participants’ heads provided visual feedback of their current CoM position in the AP direction. Two linear isolators (ULI-100, Unique Medical Co.,Ltd., Japan) were utilized to stimulate TA of both legs through 5 × 9 cm surface electrodes (PALS_®_, Axelgaard Manufacturing Co.,Ltd, USA). The electrodes were bilaterally attached at the proximal and distal ends of the muscle belly of TA, and each isolator delivered stimulation to one leg. The stimulation trains consisted of 500 μs bi-phasic, rectangular pulses, and were delivered at a frequency of 20 Hz, which was chosen to minimize muscle fatigue relative to higher stimulation frequencies [Moritani et al., 1985]. The stimulation intensity was controlled independently for each leg by a personal computer, which also collected body sway and torque data.

### Experimental protocol

#### Experiment 1

In Experiment 1, we investigated the adaptation to novel postural control dynamics. This experiment comprised three sections: (i) Estimation of the ES-torque relationship, (ii) A quiet standing task, and (iii) Evaluation of the decrease in torque production by NMES after the quiet standing task.

i. The ES-torque transfer function of the first-order system was identified from the frequency response using sinusoidally amplitude-modulated NMES pulse trains with 7 different frequencies (0.07, 0.15, 0.3, 0.75, 1.2, 1.5, and 2 Hz) [Rouhani et al., 2016] (Fig. 2a). The stimulation was delivered to TA and the resulting ankle dorsiflexion torque was measured using the custom-built torque measurement system. The maximum and minimum intensities of the sinusoidal trains were individually determined in 2.5 mA increments such that the minimum was the highest intensity that did not induce torque production, and the maximum was the highest intensity that was tolerable for each participant. The ES-torque transfer function for each leg was estimated individually to best reproduce the seven ES-torque relationships by using the MATLAB function ‘invfreqs’. Briefly, this function first calculates the vectors of polynomial coefficients of the numerator and denominator of a transfer function, *a* and *b*, by solving the following linear fitting problem using the least squares method:

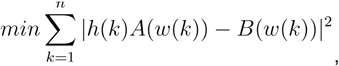

where *A*(*w*(*k*)) and *B*(*w*(*k*)) denote the Fourier transforms of the polynomials *a* and *b*, respectively, at the frequency *w*(*k*). *h*(*k*) is the frequency response, and *n* is the number of frequency points. Then, starting from the calculated *a* and *b*, the polynomial coefficients are iteratively updated by the damped Gauss–Newton method until convergence is achieved, based on the following cost function:

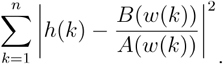 Once the ES-torque transfer function is identified, the ES pattern to generate an arbitrary ankle joint torque pattern can be obtained by plugging the torque into the inverse transfer function. The following procedure was used to test whether this idea actually worked. First, two types of ‘desired’ torque, position-dependent and velocity-dependent, were generated based on the position or velocity of the CoM AP sway from 60 seconds of quiet standing measured in advance, which were multiplied by the position- or velocity-dependent gains, respectively. These gains were individually determined such that the maximum and minimum of the ‘desired’ torque corresponded to 80% and 20% of the torque induced by NMES at the highest tolerable intensity. The NMES intensity to produce two types of the ‘desired’ torque was then estimated using the inverse ES-torque transfer function (Fig. 2d). Finally, the NMES of the estimated intensity was applied to TA, and the ‘measured’ torque was compared with the ‘desired’ torque.
ii. In the quiet standing task, the participants were instructed to stand on a force plate and maintain an upright standing posture with their eyes open. Prior to the task, the participants stood with their eyes closed for 10 seconds, and the mean CoM position in the AP direction over this period was defined as the neutral position. Figure 3a illustrates the time course of the task. The task consisted of a baseline phase, a constant-ES phase, an adaptation phase, and a washout phase. In the baseline phase, there was no NMES and no visual feedback provided. During the constant-stimulation phase, NMES was delivered at a constant intensity for 30 s without visual feedback to familiarize participants with the stimulation and to reduce excessive muscle co-contraction. The adaptation phase consisted of 12 blocks. Each block comprised a sway-dependent ES part and a probe part. The sway-dependent ES part was 30 seconds long, where the intensity of NMES was modulated based on CoM AP sway: in the position-dependent condition, the intensity was modulated based on the position of CoM AP sway, and in the velocity-dependent condition, the intensity was modulated based on the velocity of CoM AP sway. For the first and last 5 seconds of the sway-dependent ES part, visual feedback of CoM position was provided to help participants return to the neutral position. The baseline phase and each block of the adaptation phase ended with probe parts to evaluate the step response of the postural control system to a step-like NMES perturbation. The step-like NMES for the probe parts was realized by switching the NMES from low to high once the CoM was kept close to the neutral position (i.e., the velocity was below ± 0.5 cm/s and the positional deviation from the neutral position was below 0.5 cm), and the high intensity lasted for 3 seconds. Participants were asked to reduce the deviation between the current and neutral CoM position as much as possible when the visual real-time feedback was provided through the monitor in front of the participants (red lines of Fig. 3a). This ensured that the participant’s CoM position did not drift, as well as that the initial CoM position of the probe parts remained consistent. In the washout phase, the participants maintained upright standing for 60 seconds without any NMES or visual feedback, after which the final probe part was conducted. The participants underwent each condition (position-dependent and velocity-dependent) once in a random order. NMES intensity in the sway-dependent ES part was modulated to induce ankle-dorsiflexion torque represented by the following equation:

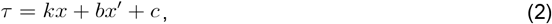

where *x* is the position of the CoM in the AP direction, *k* and *b* are the position- and velocity-dependent gains, respectively, and are a constant torque. Gain *b* was set to 0 in the position-dependent condition, while *k* was set to 0 in the velocity-dependent condition. The constants *k, b*, and *c* were determined from the participant’s body height *h* and weight *m* as follows:

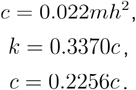 The numerical coefficients in these formulas were determined from preliminary experiments to induce sufficient body sway. The NMES intensity during the constant-ES phase was set to the same as the constant torque *c*, and the low and high intensity of the step-like NMES were set to 100% and 160% of *c*, respectively.
iii. After the quiet standing task, the ankle dorsiflexion torque was measured again when the same pre-designed velocity-dependent NMES as in section (i) was applied. The measured torque was compared to the one measured in section (i) to evaluate the degree of possible decrease in torque production due to factors such as NMES-induced muscle fatigue.

Overall, 17 males participated in Experiment 1, and 3 were excluded from the analysis due to low tolerance to NMES. Furthermore, the first three participants were excluded from the velocity-dependent condition analysis due to the use of a different velocity-dependent gain compared to the following participants. Finally, the data from 14 and 11 participants were analyzed for the position-dependent and velocity-dependent condition, respectively.

#### Experiment 2

In Experiment 2, we further investigated the influence of prolonged NMES application on torque production. Four male participants performed the same procedure as in Experiment 1 to estimate the ES-torque transfer function. We then delivered two types of NMES trains to the TA, each lasting approximately 10 minutes, separated by a 5-minute rest, while continuously measuring the resulting ankle dorsiflexion torque (Fig. 6d). Each type of NMES was pre-designed to reproduce the perturbation torque (referred to as the ‘reference’ torque) in the quiet standing task for two conditions in Experiment 1. NMES intensity of the sway-dependent ES part was determined based on the body sway during 60 seconds of quiet standing without NMES from 12 randomly selected participants in Experiment 1: Position and velocity data from 5 to 35 second segments for each participant were extracted, multiplied by the pre-defined gains, and allocated to each of 12 blocks during the adaptation phase. The gain was set to decrease exponentially during the adaptation phase to simulate the decrease in body sway associated with adaptation. The constant *k, b*, and *c* in Eq. (2) were determined similarly to Experiment 1. All participants received the position-dependent NMES first, followed by the velocity-dependent NMES.

### Data analysis

#### Torque data

For the ankle torque measurement, the force and moment during NMES application were measured by the 6-axis force sensor at a frequency of 2000 Hz and filtered by a fourth-ordered zero-lag Butterworth filter with a cutoff frequency of 10 Hz. Ankle dorsiflexion torque τwas calculated by the following equation:

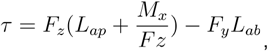

where *F*_*x*_, *F*_*z*_, and *M*_*x*_ are the forces and moment measured along each axis, *L*_*ap*_ and *L*_*ap*_ and are the distance between the ankle and the center of the force sensor along the y-axis and the z-axis, respectively (Fig. 2b). Note that this calculation does not take into account the free moment caused by friction between the foot and the measurement device. In the torque data analysis, we treated both legs separately, and thus obtained two data sets per participant.

In Experiment 2, as a primary measure to quantify muscle fatigue caused by prolonged electrical stimulation, we calculated the mean torque induced by low and high parts of the step-like NMES in each probe part. The torque exerted during 2–3 seconds after the onset of each step-like stimulation was normalized by the ‘reference’ values, and then averaged. Additionally, the reduction in torque production during the adaptation phase was quantified as the normalized mean torque for each adaptation block, defined as the ratio of the ‘measured’ mean torque to ‘reference’ mean torque.

#### Body sway data

During the quiet standing task, CoM sway in the AP direction was measured using the laser displacement sensor at a frequency of 1000 Hz and filtered by a fourth-ordered zero-lag Butterworth filter with a cutoff frequency of 10 Hz. The CoM position was then numerically differentiated to obtain the sway velocity. As behavioral measures for an adaptation to the novel postural control dynamics, we calculated the standard deviation of AP sway and the mean absolute sway velocity for each period of the baseline, adaptation, and washout phase. The first and latter half of the washout phase were defined as early and late washout phase, respectively. For the adaptation phase, only data from periods without visual feedback (6.5–25 s of each adaptation phase) were included in the analysis.

Step response to the step-like NMES was individually normalized such that the mean CoM position from 50 to 0 ms before the onset of the step-like stimulation of each step response was 0, and the peak magnitude of the step response at the baseline phase was 1. The normalized step response was then moving-averaged with a window width of three blocks. The peak magnitude was calculated as the difference between the maximum and minimum AP CoM displacement during 0 to 1.5 s after the onset. The velocity of the step response was also normalized by the maximum velocity from 0 to 1.5 s after the onset at the baseline phase. Additionally, the CoM displacement after the peak was quantified by the area under the curve (AUC) of the step response. AUC was calculated by integrating moving-averaged absolute (without normalization) step response from peak to 1 s after peak and then normalized to the value at the baseline. Stabilogram diffusion analysis (SDA) was performed on the CoM sway trajectories in the AP direction [Collins & De Luca, 1993]. The stabilogram diffusion function (SDF), which calculates mean square CoM displacement as a function of time interval Δ_*t*_, was derived from the following equation:

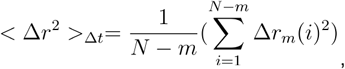

where Δ*r*_*m*_(*i*) is the difference of CoM displacement *r* between time *i* and *I + Δ*_*t*_, *N* and *m* are the number of data points in a total trajectory and Δ_*t*_, respectively, and Δ_*t*_ ranged from 0 to 10 s. The critical point coordinates *C*_Δ*t*_ and 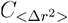 were determined as follows: *C*_Δ*t*_ was the point at which the zero-lag filtered second derivative of the SDF first became negative at Δ_*t*_ > 0.5, and 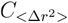 was the value of SDF at *C*Δ_*t*_. The short- and long-term regions included data ranging from 0 s to *C*Δ_*t*_ and data ranging from *C*_Δ*t*_ to 10 s, respectively. The short- and long-term diffusion coefficients *D*_*s*_ and *D*_*l*_, were calculated as half the slope of linear regression in each region. *D*_*s*_ and *D*_*l*_, were calculated as half the slope of linear regression in each region.

All sway-related parameters in the adaptation phase calculated in the aforementioned analyses, except for the ones for the step response, were smoothed by a moving average with a window width of 3 blocks due to the variable nature of the sway data. Prior to this smoothing procedure, all parameters except for SDA parameters were normalized by the values at the baseline phase.

#### Postural control simulation

In order to estimate changes in postural control gains when adapting to the new dynamics, we constructed a one-dimensional inverted-pendulum with a PID controller, following Peterka (2000) [Peterka, 2000]. The proportional and derivative gains that best reproduced each subject-averaged step responses were estimated by optimization calculations using the covariance matrix adaptation evolution strategy (integral gain was fixed at 0.02 N·m·s^-1·^deg^-1^) [Hansen et al., 2003]. Before gain estimation, the absolute step responses were first averaged across participants for all phases. For the adaptation phase only, these participant-averaged absolute step responses were further smoothed by applying a moving average with a window width of three blocks. The model height *H* and body mass *M* were set to the mean values of the participants in each condition (1.71 m, 60.6 kg for the position-dependent condition and 1.71 m, 62.4 kg for the velocity-dependent condition). The center of mass was modeled as being located at 0.55 times the body height, and the moment of inertia about the ankle joint was calculated as *I* = *MH*^2^/3.

Furthermore, we simulated AP CoM sway during the quiet standing task using the same PID controller with each set of the estimated gains. For the adaptation phase, sway was simulated under either position- or velocity-dependent perturbations, which were applied directly to the ankle joint as an external perturbation torque. For the baseline and late washout phases, we used the gains estimated from the step responses in the baseline and washout phases, respectively, whereas for the early washout phase we approximated the gains by those estimated from the last adaptation block, assuming that the control gains would not change substantially immediately after removal of the perturbation. Each simulation run lasted 40 s and was repeated 100 times for each set of estimated gains. The behavioral measures were computed from the latter half of each run, and then averaged across repetitions for each gain set. The magnitudes of the perturbation torque and step-like torque in the simulations were determined using the same procedure as in Experiment 1, except that, the position-dependent gain and velocity-dependent gain *b* in Eq. (2) was scaled to 0.8 of their estimated value to prevent divergence and oscillatory sway.

#### Statistical analysis

For Experiment 1, the accuracy of the ES-torque transfer function was evaluated based on Pearson’s correlations between ‘desired’ and ‘measured’ torque. Fisher’s *z*-transformation was used for the statistical analysis of the correlation coefficients. The reported statistics (mean and standard error) were computed by first applying this transformation to the correlation coefficients, calculating the mean and standard error of the transformed values, and then applying the inverse transformation to converge back to the original scale. To compare the correlation coefficients before and after the quiet standing task, we conducted a paired t-test on the z-transformed correlation coefficients. Paired t-tests were also used to evaluate the differences in the mean torque induced by the pre-designed velocity-dependent NMES, between before and after the quiet standing task.

We fitted a linear mixed-effects model to the behavioral measures, SDA parameters, and AUC and peak magnitude of the step response. The model contained ‘condition’ (position- and velocity-dependent), ‘time point’ and their interaction as fixed effects, and subject as a random effect. For the AUC and peak magnitude of the step response, all of the 10 time points in the adaptation phase were included in the model, whereas for the remaining variables, only the first and last time points were used. In addition to the analyses of the adaptation phase described above, we also examined differences in the washout phase for the behavioral measures and the SDA parameters by fitting analogous models with the early and late washout as ‘time point’. For each model fitting, we adopted the maximal random-effects structure that did not lead to singular fits or convergence failures. In order to evaluate the significance of the fixed effects of the models, we performed ANOVA on the fitted models with Satterthwaite’s approximation. When a significant interaction was found, post-hoc simple effects tests were performed.

For Experiment 2, a similar analysis with linear mixed-effects models was conducted on the mean torque induced by the step-like NMES and the ratio of mean torque in the adaptation phase. As fixed effects, the models for the former included ‘condition’ (position- and velocity-dependent), ‘time point’ (first and last), ‘intensity’ (low and high), and their first order interactions while the models for the latter included ‘condition’, ‘time point’ and their interaction.

For all the tests, the significance level alpha was set to 0.05. Values are reported as mean ± standard error. All statistical analyses were conducted using R version 4.3.0.

## Acknowledgments

This work was supported by KAKENHI (21H04860 and 20H05459), TRIBAWL Co., Ltd., and MTG Co., Ltd. to D.N., and KAKENHI (19K20067) to S.H.

## Author contributions

Conceptualization and methodology, R.M., S.H., and D.N.; formal analysis, software, and investigation, R.M. and S.H.; visualization, R.M. and D.N.; resources, D.N.; writing—original draft, R.M.; writing—review and editing, R.M., S.H., and D.N.; funding acquisition, D.N.; supervision, D.N. All authors have read and agreed to the published version of the article.

## Figure

**Supplementary Figure 1.**
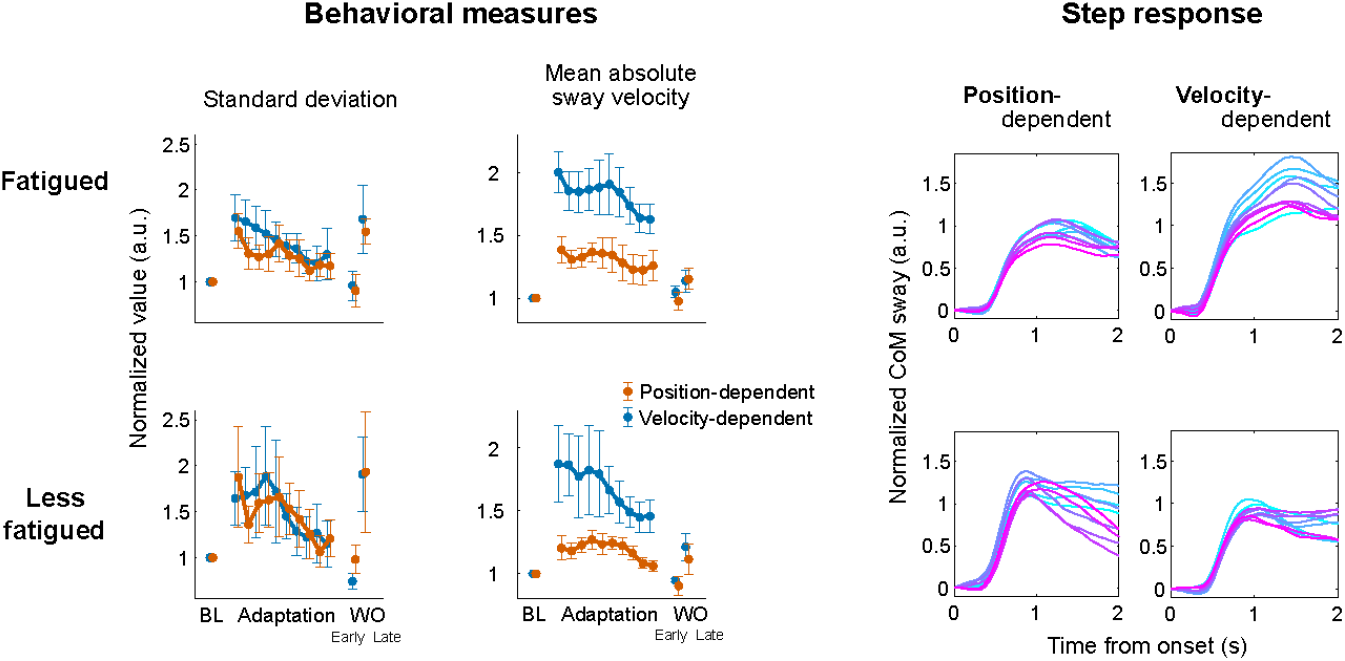
Subgroup analysis for the behavioral measures (left side) and step response (right). Participants were divided into two groups based on the degree of reduction in mean ‘measured’ torque after the quiet standing task. Participants exhibiting a reduction of greater than 40% were assigned to the fatigued group (top; n = 8 for position-dependent and n = 7 for velocity-dependent), and the remaining participants were assigned to the less-fatigued group (bottom; n = 6 for position-dependent and n = 4 for velocity-dependent). Error bars correspond to SEM across participants. Statistical comparisons were not performed due to the small sample size.

